# Diversity of GABAergic interneurons and diversification of communication modules in cortical networks

**DOI:** 10.1101/490797

**Authors:** Z. Josh Huang, Anirban Paul

## Abstract

The phenotypic diversity of cortical GABAergic neurons is likely necessary for their functional versatility in shaping exquisite spatiotemporal dynamics of circuit operations underlying cognitive processes. Deciphering the logic of this diversity requires overcoming the technical challenge of quantitative and comprehensive analysis of multi-modal cell features as well as the conceptual challenge of formulating a framework of neuronal identity that reflects biological mechanisms and principles. In the past few years, advances in high-throughput single cell analyses began to generate unprecedented datasets on interneuron transcriptomes, morphology and electrophysiology that drive their classification. Recent studies suggest that cardinal interneuron types can be defined by their synaptic communication properties encoded in key transcriptional signatures – a conceptual framework that integrates across phenotypic features and captures neuronal input-output properties elemental to circuit operation. This definition may further facilitate understanding the appropriate granularity of neuron types toward building a biologically-grounded and operationally useful interneuron taxonomy.

## Introduction

Understanding the biological basis of neuronal diversity is necessary for deciphering neural circuit organization and function. Across the mammalian central nervous system, GABAergic neurons of the neocortex and hippocampus are among the most intensely studied cell classes. Spectacularly diverse GABAergic neurons appear to represent an elaborate division of labor in deploying a rich repertoire of inhibitory control mechanisms to shape highly nuanced spatiotemporal dynamics of cortical circuit computation (*1*, *2*). Since the invention of Golgi stain that unveiled the elaborate shapes of these “short axon cells” by Ramon y Cajal, our appreciation of the anatomical, physiological and molecular diversity of GABAergic neuron has been driven by waves of technological advances in microscopy, neurochemistry, electrophysiology and molecular biology. Decades of classic studies have led to seminal discoveries of the stunning specificity of interneuron phenotypic features in terms of their synaptic connectivity (e.g., selectivity in synaptic partners and subcellular compartments), physiological characteristics (e.g., intrinsic and especially synaptic properties), and functional properties in circuit operations (e.g., temporal integration and network oscillation) (*1-4*). These studies have also revealed immense, and often seemingly intractable, phenotypic variations along multiple axes that defy any simple classification scheme (*5*). As a result, the issue of interneuron diversity remains contentious (*6*). There have been continued debates on how interneuron “types” should be defined and classified, and what is the scope of their diversity (i.e. how many types and at what granularity). Indeed, many important problems in circuit neuroscience can often be traced to the ambiguity of “neuron types” in the system under investigation (*7*). At the core of these debates is the issue of whether neuronal identity and classification can be grounded on underlying biological mechanisms and principles, or whether they are destined to remain arbitrary and operational.

Given the immense complexity of the cortical circuits in which GABAergic interneurons are embedded, our current knowledge gap and incomplete understanding are not surprising. The multi-modal and multi-dimensional phenotypes of nerve cells are extraordinarily difficult to describe, measure and comprehend. These include not only morphology, connectivity patterns, physiological properties, and gene expression profiles but also developmental history and ultimately circuit and behavioral function. Despite major advances in past decades, until recently investigators remain technically underpowered, to a large extent, to meet these challenges. For example, sparse and partial (instead of systematic and complete) reconstructions of individual cells preclude truly quantitative analysis of cell morphology. Most electrophysiological recordings sample limited and often biased cell populations and only a fraction of the physiological parameter space that neurons operate in, especially considering the diverse input-output transformation properties among cell types. Serendipitously identified molecular markers represent a minute fraction of gene expression profiles. Thus overall these studies remain severely limited in resolution, robustness (e.g. reproducibility across investigators) and comprehensiveness.

The conceptual obstacle to understanding neuronal diversity manifests at multiple levels. First, cell features often display substantial multi-dimensional variations that appear continuous as well as discrete (*8*, *9*). As we often cannot distinguish between biologically meaningful variations versus stochastic or technical variations, it is difficult and often arbitrary to set boundaries for cell clustering by adjusting algorithmic parameters. Second, the inherently multi-modal nature of cell phenotypes requires integrative classification across modalities, but it is not clear whether different cell features indeed co-cluster, raising the issue of whether congruent multi-modal classification is achievable. Third, it seems intuitive that neurons should be viewed as members of a type if they serve a common function, and cell typing is only useful if it ultimately helps understand such function(s). In practice, however, cell “function” is not readily definable and emerges only at the circuit level, which is difficult to study and is often indirectly linked to behavioral performance. At the core of these challenges is the lack of an overarching framework of neuron type identity that integrates and explains multi-modal variations relevant to circuit operation and is grounded on biological mechanisms and principles.

In the past few years, spectacular advances in single cell analysis are finally crossing several technical thresholds and begin to generate high resolution, quantitative and comprehensive datasets in single neuron transcriptomes, morphology and electrophysiology. Innovative statistical and computational analyses allow clustering and typing along each modality and drive efforts to integrate across modalities. Here we review progress in large scale single cell RNA sequencing (scRNAseq) that unveiled a working draft of a comprehensive transcriptomic interneuron taxonomy, and the related morphological and electrophysiological datasets that may contribute to multi-modal classification. We highlight an emerging framework of neuronal identity, which defines cardinal interneuron types as canonical neural communication modules with characteristic input-output transformation properties encoded in transcriptional signatures of key gene families. We discuss outstanding challenges and opportunities for future progress.

### High throughput single cell analyses toward classification and taxonomy

#### Single cell transcriptomics

To a significant extent, cell phenotypes and properties derive from their patterns of gene expression. Since the 1980s, various molecular markers (e.g. calcium binding proteins and neuropeptides) have been used to parse GABAergic subpopulations (e.g. (*10*, *11*)), but these serendipitous markers are limited in distinguishing cell types. More recently, transcription profiles of GABA subpopulations have been characterized through microarray or RNA sequencing (*12*), but these studies using bulk cell populations have inherent limitations in resolving heterogeneity, which manifests in individual cells. Only in the past three years, large scale scRNAseq has finally crossed the threshold for molecular analysis of neuronal diversity (*13*, *14*). In the mouse primary visual cortex, a pilot study of ∼1600 cortical cells combined with statistical clustering identified 49 cell types, including 23 GABAergic transcriptomic types (*15*). A subsequent large scale study of ∼23,800 cells incorporating multiple transgenic lines suggests 113 cortical cell types; these include 61 GABAergic types that are conserved between visual and front cortex and 56 glutamatergic types that are mostly distinct between these two areas (**Figure 1**)(*16*). This landmark dataset represents the most comprehensive survey of cortical transcriptional types, establishing a scaffold for transcriptome-based classification and a work draft of cortical cell taxonomy, including GABAergic interneuron taxonomy. The major subclasses, types and their hierarchical relationships are largely consistent with previous studies (*4*, *11*, *17*). Some of the finer types and branches (e.g. Sst-Chod1, Sst-Calb2-Pdlim5, Sst-Chrna2-Glra3) are also consistent with previous studies (**Figure 1**)(*18-20*). Certain clusters may correspond to newly discovered types; for example, one of the four clusters defined by VIP and Chat expression might correspond to a type that directly excites layer 1 interneurons and layer 2/3 pyramidal neurons (*21*).

**Figure 1.**
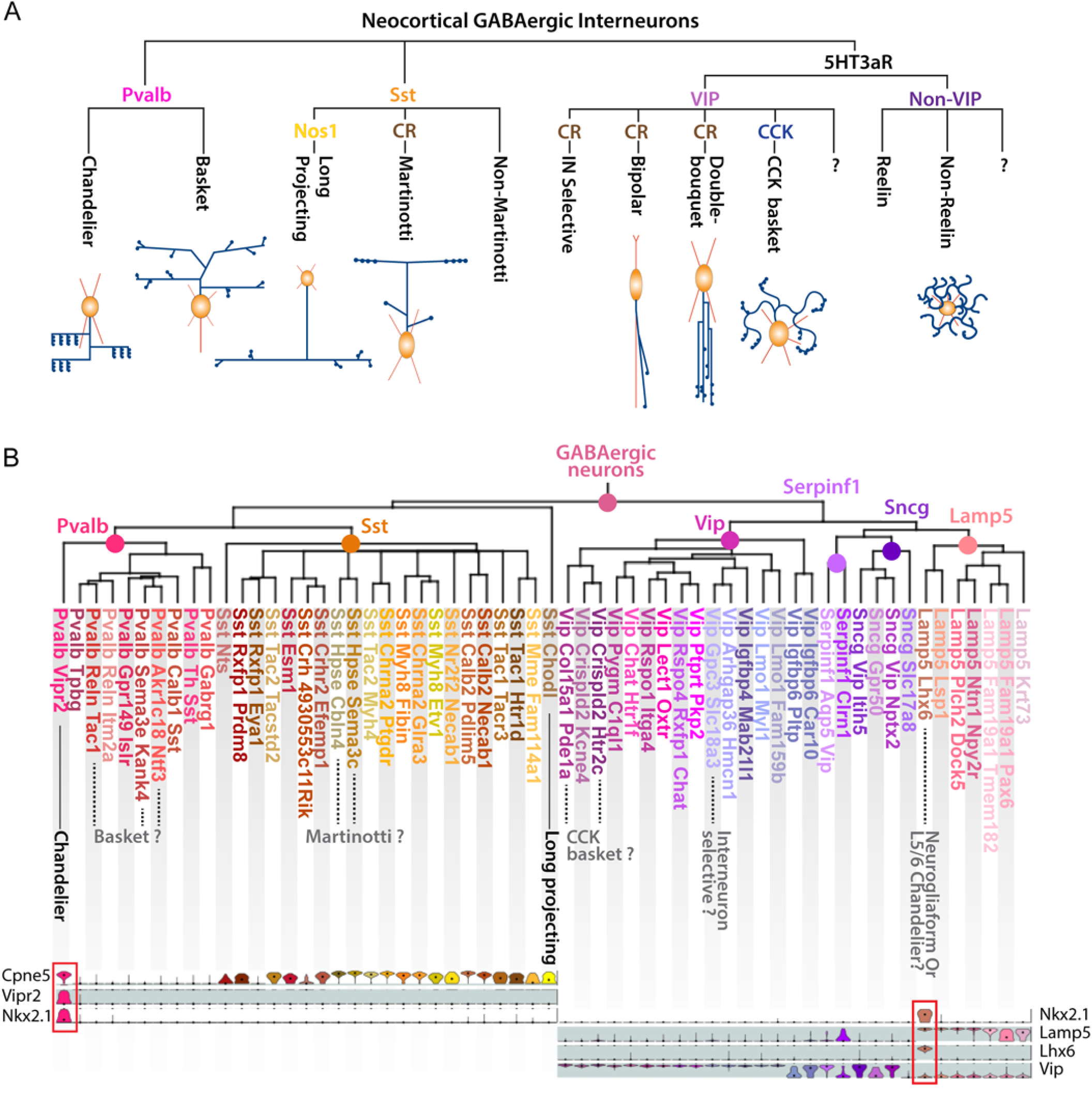
A work draft taxonomy of transcriptomic neuron types of the cortical GABAergic system (adapted from (*16*). A) Major GABAergic subclasses and cell types recognized by classic anatomic, physiological, molecular and developmental studies. B) Current taxonomy of transcriptomic types in the primary visual and anteriolateral cortex. Note that the major types and branches are consistent with classic studies. Only a few (bold) of the 61 “atomic types” are firmly correlated with the anatomically and physiologically defined types. A major discrepancy is the Lamp5-Lhx6 type, which is clustered as CGE derived NGF cells (*16*) but was recognized as MGE derived CHC (*20*).

However, the taxonomy of cortical GABAergic neurons is not settled. The ∼61 “atomic” GABAergic transcriptomic types at the base of the hierarchy (*16*) are likely fluidic and may well be modified by future scRNAseq and orthogonal datasets. Currently, the choices of statistical algorithms remain largely arbitrary – the stringency and criteria for cluster definition are up to individual researchers (i.e., how many genes and how differentially expressed they need to be between for two potentially distinct types to be annotated as such) (*22*). For example, the number of Sst transcriptional clusters can vary by 2 fold depending on the stringency of clustering (30 when less stringent and 15 when more stringent) (*16*). These results accentuate the issue of the extent that “neuron types” are statistical or biological.

There are also a number of results that are inconsistent with previous studies. For example, decades of developmental studies have established a clear division of transcription programs between the medial and caudal ganglionic eminence derived interneurons (*3*, *23-25*). In particular, the *Nkx2*.*1* and *Lhx6* transcription cascade drives the MGE but not the CGE clade (*23*, *26*, *27*). Interestingly and surprisingly, a group of *Lamp5* subclass interneurons with prominent *Lhx6* and *Nkx2*.*1* expression was classified as CGE derived putative neurogliaform cells (NGFC) (*16*). Notably, this Lamp5-Lhx6 type was identified as transcriptionally similar to the CHC2 type, which was considered L5/6 chandelier cells that were fate mapped from Nkx2.1 progenitors in the MGE clade (*20*). It will be important to resolve this major discrepancy as it is rooted at the top level hierarchy of interneuron classification. One possibility is CHC2 is in fact NGFC, and *Nkx2*.*1* and *Lhx6* are indeed not strictly restricted to the MGE lineage and somehow also generate certain CGE cell types. This would be a novel discovery casting doubt on the stringency of core transcription programs separating the fundamental MGE and CGE lineages. Another possibility is that Lamp5-Lhx6 is in fact MGE derived deep layer chandelier cells. In this scenario, non-essential transcriptional features (e.g. derived from technical or tangential factors) could have over taken key biologically meaningful signals and misled the clustering algorithm to assign them as NGFC in the CGE clade. Consistent with this possibility, *Lamp5* is also expressed in L2 ChCs ((*20*); online expression dataset), suggesting that *Lamp5* is not a CGE specific interneuron marker as proposed (*16*). Resolving this discrepancy will not only clarify the distinction between two well characterized bona fide types (NGF vs CHC) but also may reveal the power as well as glitches in statistical clustering. For example, it might be necessary to supervise clustering algorithms so that foundational core transcriptional factors (e.g. *Nkx2*.*1, Lhx6*) have higher voting power in clustering cell identity and relationship than other genes.

The current fluidic nature of transcriptomic clustering is at least in part due to our incomplete understanding of the biological basis of neuron type identity and granularity, and the paucity of “ground truth” knowledge. Such understanding likely requires integrating multi-modal cell features, recognizing biological meaningful variations in each features, and discovering their mechanistic basis.

#### Single cell epigenomics

Transcriptomes in individual cells are transcriptional outputs of their epigenomes, which are customized from a singular genome largely through developmental genetic programming. In addition to scRNAseq, technical advances now allow the delineation of genome wide DNA methylation patterns (*28*) and chromatin accessibility (ATACseq (*29*) in single cells. Although yet to reach the resolution and scale of scRNAseq, these single cell epigenomics methods begin to resolve major cortical cell classes and subclasses (e.g. GLU, GABA PV, Sst neurons and glia cells etc). With further technical improvement, an integrated analysis of single cell transcriptomic and epigenetic datasets (*30*) will not only facilitate molecular cell typing but also may provide deeper insight into the underlying gene regulatory basis.

#### Single cell morphology

It seems intuitive to describe neuron types based on their morphology. However, the vast diversity and seemingly endless variations of neuronal shapes present major challenges in morphological tracing and analysis. A century after the description of “short axon cells”, it has remained difficult to achieve rigorous, quantitative and scalable analysis of interneuron morphology toward their classification (*6*). Quantitative single neuron anatomy requires overcoming four technical hurdles. The first is labeling - to systematically, reliably, sparsely and completely label specific sets of individual neurons. The second is imaging - to achieve axon resolution large volume imaging brain-wide. The third is reconstruction - to convert large image stacks into digital datasets of single neuron morphology. The fourth is analysis - to register neuronal morphology with appropriate spatial coordinate framework and to extract, quantify and classify biologically relevant attributes (e.g. those related to neural connectivity). Most studies in past decades using traditional methods can only recover partial interneuron morphologies that are not well registered to a common spatial coordinate framework. These limitations make it difficult to achieve quantitative analysis, distinguish random or technical variation versus reliable and meaningful features, and compare across investigators. A tour de force large scale study in juvenile rat brain slices digitally reconstructed 1009 cortical neurons (*17*). Statistical analysis combined with literature mining and expert annotation classified these into 55 morphological types, including ∼40 interneuron types with their associated laminar locations. Another study of the adult mouse visual cortex took advantage of a large set of transgenic lines and reconstructed 199 spiny and 173 aspiny neurons (*31*). Applying unsupervised hierarchical clustering, they identified 14 spiny morphological types (m-type) and 21 aspiny m-types. Together these two studies represent the most comprehensive morpho-physiological analysis of cortical interneurons to date. Despite these progresses, brain slice preparations present inherent limitations for complete morphological reconstruction, precise spatial registration and truly quantitative analysis.

The recent integration of genetic labeling and axon resolution, large volume imaging has begun to overcome major technical hurdles of single neuron anatomy in rodent brains. In one platform, high-speed two-photon microscope is integrated with sparse viral labeling of neurons and computational tools for large scale image analysis (*32*). In another platform, dual-color fluorescence Micro-Optical Sectioning Tomography (dfMOST) allows axon resolution imaging and cell resolution spatial registration of genetically labeled single neurons (*33*). Combined with genetic labeling, a recent study achieved complete reconstruction of over 60 chandelier cells (ChCs) in mouse prefrontal, motor and somatosensory cortex. Analyses of their laminar position, dendrite and axon arbor distribution suggest multiple ChC subtypes are likely distinguished by different input-output connectivity (*34*). These technical advances signal the rise of high resolution, quantitative and scalable single neuron anatomy in the mouse brain. Currently, a major bottleneck is morphological reconstruction, which is mostly achieved by manual procedures that are particularly labor intensive. Innovations in more automated reconstruction and registration will be necessary to generate high throughput and comprehensive dataset. In terms of analysis, conceptual as well as technical questions remain: how to distinguish stochastic variations versus biologically meaningful features? Can morphology truly inform cell typing? As morphology is a proxy of connectivity, how should it be interpreted in the context of connectivity (*35*)?

#### Electrophysiology

The electrophysiological properties of neurons are proximal to their roles in circuit operation yet are even more difficult to measure comprehensively and quantitatively at scale, especially in vivo. Different neuron types likely mediate distinct sets of input-output transformations supported by their intrinsic, synaptic and network properties which span orders of spatiotemporal scales from dendritic spines to axon terminals and from sub-milliseconds to seconds; these are further influenced by brain state and behavior. Largely due to technical limitations, only a small set of physiological properties can be routinely measured, the most common being intrinsic properties at cell soma and a limited set of synaptic properties, predominantly in brain slices in vitro. For example, Markram et al 2015 applied a standardized battery of stimulation protocols to >3,900 neurons throughout rat cortical layers and analyzed their responses, identifying 11 electrophysiological types (e-types) (10 inhibitory types and 1 excitatory type). Using a similar approach to intrinsic properties and aided by a set of transgenic driver lines, Gouwen et al 2018 classified 6 excitatory e-types and 11 inhibitory e-types. While these datasets are very valuable, intrinsic responses measured at cell soma in vitro provide a very limited glimpse of the highly sophisticated spatiotemporal operations of neurons in their network niche.

Measuring synaptic properties requires multiple simultaneous patch clamp recordings that are technically demanding and even more difficult to scale. An impressive profiling of cortical cell types using octuplet whole cell recording in adult mouse cortex mapped connectivity between over 11,000 pairs of identified neurons (*36*). They categorized 15 types of interneurons, each exhibiting a characteristic pattern of connectivity with other interneuron types and pyramidal neurons. However, as each neuron receives inputs from and transmit outputs to multiple pre- and post-synaptic neurons, respectively, a more comprehensive profiling of synaptic signaling is prohibitive even with octuplet patch recording by expert physiologists.

Regarding network properties, the firing pattern of hippocampal interneuron types during various in vivo network oscillations are highly characteristic (*37*, *38*), but because cortical network oscillations are less discernable, similar measurements are not yet feasible for cortical interneurons. Therefore, current measurements of the electrophysiological properties provide a severely limited glimpse into their sophisticated and likely characteristic spatiotemporal and network operations. As most common datasets on intrinsic properties in vitro may not reflect the essence of neuronal physiological operations, it is perhaps not surprising that they have not been particularly informative in neuronal classification. Proper distinction of physiological neuron types may require distilling overarching functionally relevant features, such as integrating multiple methods to measure the input-output properties in the context of circuit operations. Such measurements at large scale may require the innovation of novel technologies.

### Correspondence among multi-modal datasets

With the broad application of scRNAseq, it is now possible to begin to correlate transcriptomic, morphological, and physiological datasets toward a multi-modal cell classification scheme. One approach is Patch-seq, which combines whole-cell patch clamp recordings, scRNAseq and morphological characterization (*39*, *40*). Another approach is to integrate multi-modal measurements of the same neuronal subpopulation labeled by common mouse driver lines. In an impressive large scale study, Gouwens et al 2018 analyzed the intrinsic properties of 1851 neurons (885 spiny and 966 sparsely spiny) and reconstructed the partial morphology of 372 (199 spiny and 173 aspiny) neurons in adult mouse visual cortical slices. These cells derived from a comprehensive set of 28 Cre driver lines to ensure broad coverage, to enable selective targeting of rare subpopulations, and to link these datasets to single cell transcriptomic datasets of the same subpopulations (*16*). Among the 17 e-types and 35 morphological types (m-types) identified, there is an overall trend of correspondence between groups of e-types and m-types. However, with rare exceptions (e.g. thick tufted cells in L5 – spiny 7 m-type with Exc_3 e-type), there were no clear correspondences between e- and m-types. To examine the relationship between transcriptomic types (t-types) to e- and m-types, Gouwens et al took advantage of several driver lines that each correspond to a small number of t-types and the same lines were used to characterize e- and m-types. The best example of multi-modal correspondence is the L5/6 Sst Chodl long projection GABA neurons that were captured by Nos1/Sst intersection (*19*, *20*). In general, however, the correspondence among t-, e-, m-types are coarse and weak. Thus the extent to which the 61 t-types correspond to the 11 e-types and 21 m-types are currently unclear.

Taking these results on face value, one might tend to conclude that there is unlikely tight correspondence among the transcriptomic, morphological and electrophysiological features of GABAergic neurons. However, such a conclusion is premature for several reasons. Given the fact that intrinsic properties measured by arbitrary stimulation parameters in vitro is a poor representation of the overall electrophysiological features and the limitation of morphological analysis of partially reconstructed neurons from brain slice preparation, the current relatively weak correspondence between e and m types is not surprising. Further, as morphological and physiological measurements are at a far lower resolution than transcriptomic measurements (thousands of dimensions related to the number of mRNAs expressed in a cell), the weak correspondence between the 61 t-types to 11 e-and 21 m-types are also not unexpected. It is possible that the current ambiguity and disparity in cell clustering across modalities reflect the lack of a conceptual framework that captures the essence of neuron type and distill the core overarching properties that can encapsulate and possibly extend beyond the traditionally described cell phenotypes.

### GABAergic neuron type is defined by communication properties rooted in transcriptional signatures

#### Transcription profiles of key gene families customize cellular modules that shape neuronal input-output properties

Despite progress in large scale single cell transcriptomics, the problem of neuronal diversity and classification is unlikely to be solved without understanding the nature of neuronal identity and the fundamental principles guiding the organization of cell types. Significant progress has been made by analyzing high resolution single cell transcriptomes leveraging the deep, even though incomplete, knowledge in GABAergic neuron anatomy and physiology (*20*). This study differs from other single cell transcriptomic studies in three significant ways (Figure 2). The first is experimental design. Instead of aiming to discover diversity and classify neurons using unsupervised statistical clustering of transcriptomes from unbiased or relatively broad populations, Paul et al focused on understanding the molecular basis of neuronal identity by analyzing the transcriptomes of 6 phenotypically-characterized populations (PCP) of GABAergic cardinal types (each comprising ∼100 individual cells) labeled by combinatorial genetic targeting. The phenotypic knowledge in these PCPs derive from decades of anatomical, physiological (including synaptic and network properties) and developmental studies, which provide a solid basis for the interpretation of and correspondence to the transcriptomic dataset. The second is scRNAseq method. Using a modified version of CEL-seq with two-round in vitro transcription combined with unique molecular identifiers (UMI), Paul et al were able to detect ∼9,000 genes per single cell) and achieve mRNA counting in single cells. This method thus achieves substantially higher resolution of transcription profiling compared with Drop-Seq and 10x Genomics platforms and a more quantitative mRNA measurements than Smart-Seq which does not incorporate UMIs. The third is analysis. Instead of using unsupervised algorithms to cluster cells based on overall transcription profiles, this study aimed to extract key transcription features (e.g. gene families) that contribute to, explain and predict cell properties. The analysis strategy is based on the premise that cell phenotypes emerge from operations of macromolecular machineries (i.e. cellular modules) comprising interacting protein components (*41*), each is often implemented as one of multiple variants encoded by a gene family. Thus differential expression of gene family members with characteristic biochemical and biophysical properties may confer tailored features to cellular machines thereby customizing cell phenotypes that together characterize a cell type (Figure 2).

**Figure 2.**
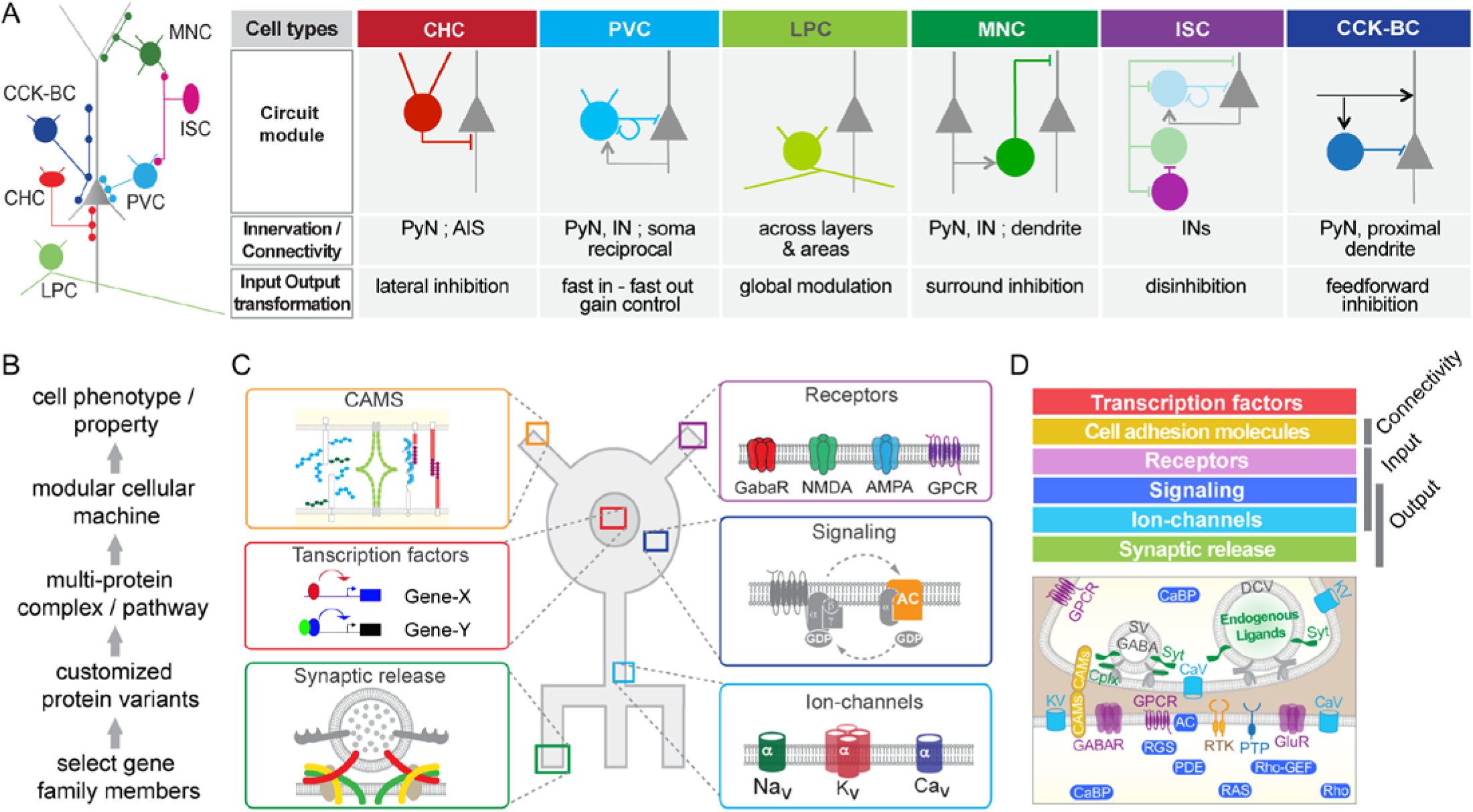
Transcriptional signatures of synaptic communication delineate cardinal GABAergic neuron types. A) Six cardinal types are distinguished by their characteristic innervation of cellular and subcellular targets. LPC-long projection cells, CHC-chandelier cells, BC-basket cells, ISC-interneuron selective cells, MNC-Martinotti cells. Where data available, these cardinal types manifest distinct input-output connectivity patterns and further display distinct intrinsic, synaptic and network properties indicative of mediating specific forms of input-output transformation. PyN-pyramidal neuron; AIS-axon initial segment; IN-interneuron. B) Cell phenotypes and properties emerge from the workings of macromolecular machines (cellular modules) consisting of multi-protein complexes; many components are implemented as variants with different biochemical and biophysical properties encoded by members of a gene family, which customize cellular modules and cell type specific properties. C) Examples of cellular modules that shape neuronal connectivity, synaptic transmission, electrical signaling, intracellular signal transduction and gene transcription. D) Six categories of gene families shape a set of membrane proximal molecular machines that customize the input-output connectivity and transformation properties of different GABAergic cardinal types.

Using the known PCP identities of ∼600 single cell transcriptomes as an *assay*, Paul et al designed a supervised machine learning-based algorithm (MetaNeighbour; (*42*)) to *screen* through the ∼620 HGNC (Human Genome Nomenclature Committee) gene families for those whose differential single cell expression reliably distinguish these PCPs. Remarkably, only ∼40 gene families, mostly implicated in regulating synaptic communication, best distinguished the PCPs. These gene families constitute 6 functional categories that include cell-adhesion molecules, neurotransmitter and modulator receptors, ion channels, regulatory components of membrane-proximal signaling pathways, neuropeptides and vesicular release components, and transcription factors. Combinatorial expression of select gene family members across these categories shapes a molecular scaffold along the cell membrane that contribute to pre- and post-synaptic properties, i.e. input-output (I/O) characteristics (Figure 2). These findings suggest that a cardinal neuron type is a neural communication module that is rooted in transcriptional programs orchestrating functionally congruent expression across a key set of gene families to customize the patterns and properties of I/O transformation. This is consistent with an independent proposal to define projection neuron types based on input-output relationships (*43*).

#### An overarching and mechanistic framework of neuronal identity

The definition of neuron type as a communication module rooted in its transcriptional program represents a higher level abstraction that integrates multi-modal cell phenotypes, provides a framework for understanding neuronal diversity, and may facilitate cell type classification based on biological principles beyond statistical algorithms and operational management (Figure 3). The two fundamental features of a communication module are its connectivity (i.e. with whom to communicate) and input-output transformation (i.e. how to communicate), which encapsulate as well as extend the more conventional descriptions of cell features.

**Figure 3.**
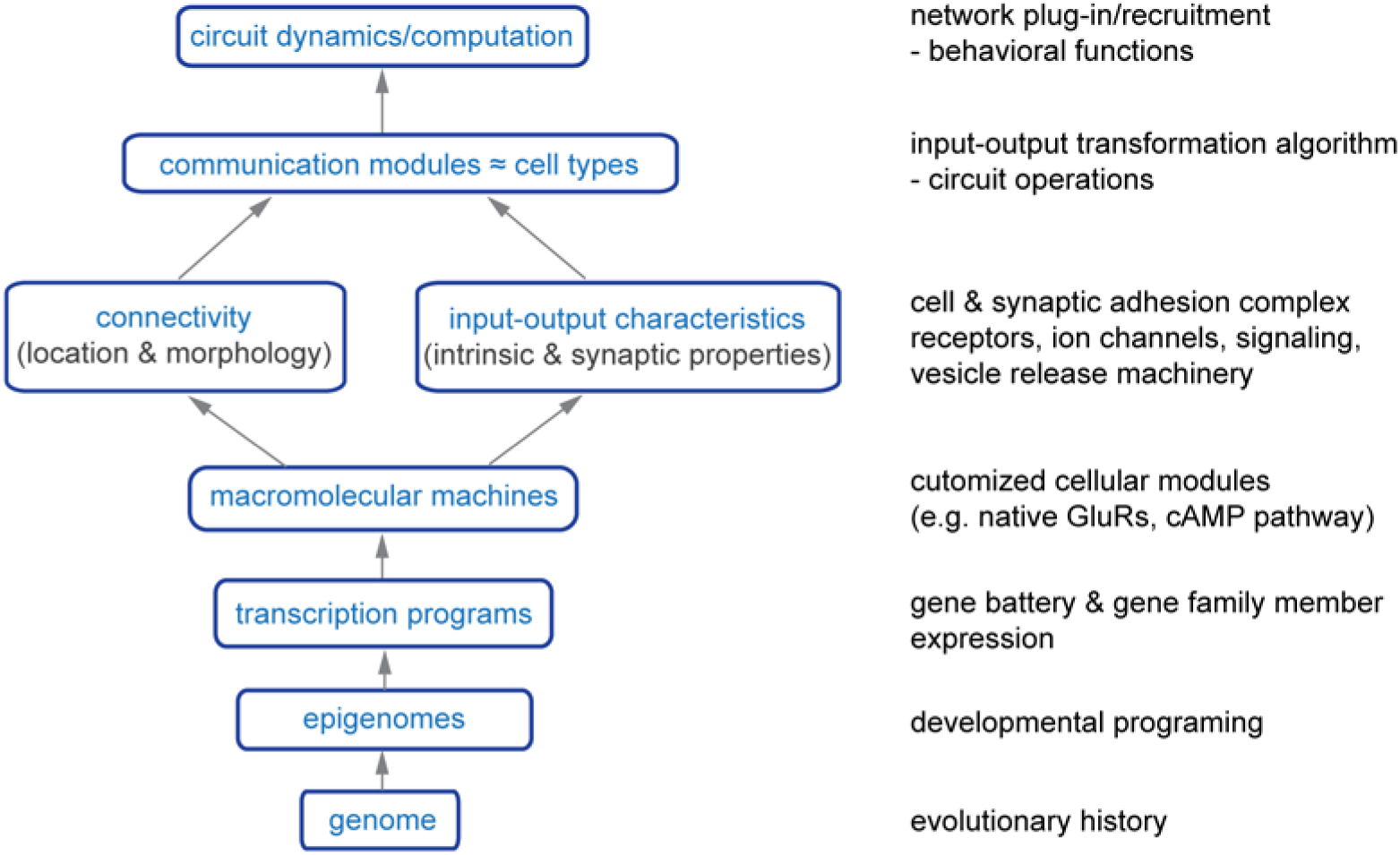
A neuron type can be defined as a canonical neural communication module mediating characteristic input-output transformation and is encoded by key transcription signatures that customize a congruent set of cellular machines. This scheme may integrate anatomical, physiological, functional, and developmental genetic features that together define neuronal cell types. It emphasizes cardinal neuron types that are reliably generated in each members of the species, build species-stereotyped circuit scaffolds and templates and are likely rooted in the genome; not shown is the notion that these cardinal types are likely further diversified and customized, e.g. by neuronal activity and experience, to shape circuit elements characteristic to individual organisms. Features in gray font are commonly measured cell properties but are reflections of core features of cell types. See text for detailed description.

In terms of anatomical features, for example, although location and morphology have been prime descriptors of neurons, they reflect and serve the more fundamental purpose of achieving proper connectivity. Thus seeming intractable morphological variability may belie the co-variation of pre- and post-synaptic neurites that preserves the same connectivity pattern (Seung and Sümbül, 2014). Indeed, morphological types can be reliably identified from dense connectomes by computational algorithms (*44*, *45*). In terms of physiological features, the intrinsic, synaptic and network properties of a neuron type likely all serve the overarching goal of transforming information contents embedded in its synaptic inputs (e.g., transmitter and modulator types, strength, and spatiotemporal dynamics) to appropriate pattern and dynamics of synaptic outputs, which are often characterized by cell intrinsic style of neurochemical release (e.g., vesicle contents and release speed, dynamics, and plasticity). A functional definition of neuron type seems intuitive and the most relevant to nervous system operation (*3*, *43*), but neuronal functions manifest at multiple levels, carry different meanings to different investigators and thus seem unsuited to define cell types. A key feature of the communication module concept is that it seamlessly incorporates the functional definition of neuron types across levels. Neuronal communication modules equipped with specific input-output transformation algorithms carry out a specific set of definable circuit operation (e.g. feedforward inhibition, lateral inhibition, disinhibition, gain control, gamma oscillation) – their proximal level function. Plugging-in and task-dependent recruitment of these modules into larger brain networks then engage their systems level information processing and computation that contribute to their distal behavioral and cognitive functions (*46*, *47*). Thus the communication module concept incorporates the functional definition of neuron types by clarifying and integrating different levels of functional explanations. It is likely that cardinal cell identity is already instilled at the postmitotic stage during development (*3*, *48-50*), and its subsequent unfolding and expression are regulated by differentiation signals such as cell interactions and neural activity. Importantly, key communication properties are shaped by specialized cellular modules encoded in transcriptional programs. Although transcription profile is influenced by cellular milieu including neural activity (*51*), core transcription programs are grounded in cellular epigenomes customized primarily through developmental programming of the genome. Therefore, the definition of neuron type as communication module encoded in transcriptional signatures of I/O properties coherently integrates anatomical, physiological, functional, and developmental genetic features that together define neuron types (Figure 3).

An intriguing corollary is that communication is inherently a cell extrinsic feature tightly linked to pre- and postsynaptic cells, yet it is rooted in cell intrinsic transcription programs. This suggests that a cardinal cell type may be endowed with genetic information to cell autonomously seek other appropriate cell types for establishing characteristic connectivity patterns (*50*) as a central step of implementing and expressing their core identity. It is thus possible that cell intrinsic transcription programs in diverse cardinal neuron types, which derive from their epigenomes, is a key intermediate step through which the genome ultimately orchestrates the self-assembly of canonical features of neural circuit scaffold.

In this context, it is perhaps not surprising that there is a relatively poor correlation between currently described morphological and electrophysiological types, as morphology is an indirect measurement and reflection of connectivity whereas electrophysiological properties at cell somata is only one of many contributors of I/O transformation. It is possible that key connectivity and I/O features are in fact highly correlated once they can be measured at high resolution. It is also possible that certain morphological features are more tightly linked to connectivity (e.g. chandelier cell axon arbors; (*34*) and certain intrinsic and synaptic features are better linked to I/O (e.g. fast in-fast out in fast-spiking basket cells; (*52*)). Once identified, their tighter correlations in specific subpopulations may begin to emerge.

#### Communication module as a testable hypothesis

The communication module definition of neuron type should be viewed as a testable hypothesis, with the prediction that input-output transformation properties correlate with transcriptional signatures that together define neuron types. Recent development of tools that combine genetic targeting (*19*, *53*), antero-, retro- and trans-synaptic viral tracing (TRIO etc; (*54*)), and optogenetic mapping are well poised to facilitate the discovery of the I/O streams of neuronal subpopulations (*43*, *55*). In parallel, dense connectomics datasets by volume electron microscopy may reveal the complete connectivity of individual neurons and allow investigators to discern key patterns that define cardinal anatomic types. Current efforts on EM reconstruction of a cortical column may reveal the local connectomes of interneurons and begin to allow investigators to examine the relationship between connectivity and cell identity (*56*). Larger volume EM connectomes will be necessary to fully examine the I/O streams of cortical interneurons. In terms of I/O transformation, currently there is no single method to measure this higher level neuronal operation. Instead, different components such as synaptic inputs and integration, spike initiation and propagation, and synaptic release are measured separately only in a few abundant and accessible cell populations (e.g. PV fast-spiking basket cells, (*52*)). On the other hand, technical development increasingly enables subpopulation-targeted and wiring-dependent optogenetic control, activity readout and perturbations, which allow probing I/O transformation grounded on cellular resolution anatomical pathways (*43*). Despite these progresses, large scale measurement of I/O transformation in neuronal subpopulations in vivo remains a significant challenge and may require entirely new technologies. Importantly, I/O transformation properties appear to be encrypted in key transcription signatures (*20*). Proper analysis of high resolution transcriptomes may guide the characterization and validation of their I/O patterns and facilitate the discovery of neuron types (i.e. beyond statistical transcriptional types). Recent advances in spatial transcriptomics and their increasing integration with modern anatomical and imaging tools (*57-59*) may facilitate correlating I/O properties with transcription signatures. Applying this strategy to a modest set of cardinal types, e.g. glutamatergic pyramidal neurons, seems feasible. Once the correspondence between key transcription signatures and I/O properties is established, scalable high-resolution transcriptome analysis may guide the large scale discovery of cardinal neuron types.

### How to define the appropriate granularity of interneuron types?

In addition to understanding the nature of neuronal identity, it is necessary to recognize and define the appropriate granularity of cell type boundaries in order to achieve meaningful classification and taxonomy (*9*). The question of how many cell types a given brain region contains is frequently asked, but this question will not have a clear answer until the investigator can determine how finely and firmly cell types should be distinguished that is appropriate for understanding circuit function. Inherently related to the issue of neuron type identity, the question of granularity presents similar technical and conceptual challenges at a finer resolution.

Even neurons recognized as members of a given subclass or type often further manifest significant multi-modal variabilities that appear discrete as well as continuous. It is unclear as to when and whether investigators should continue to finely cluster cells along each modality to define “subtypes”. For example, single cell transcriptome datasets manifest discrete as well as continuous variations (*8*, *15*). A major question is what variations are biologically meaningful, and which ones are not (e.g. technical or stochastic). Similarly, morphological variations of individual neurons of the same class may seem continuous and intractable (*6*), but these may belie the more specific and discrete connectivity patterns when both pre- and post-synaptic elements are considered from dense connectomics datasets (*60*, *61*). Along the same vein, the significance of the variation of intrinsic physiological properties may only be interpretable in the context of their impact on I/O transformation. Although some argue for an emphasis on discrete variables (*9*), others suggest continua as a necessary description of cell diversity (*8*). These debates uncover a deeper conceptual issue: what level of granularity of cell type definition is necessary for understanding circuit function?

In this context, the communication module concept of neuron types may provide a basis and guideline for defining its granularity, with the prospect of linking variations of key cell features to their impact on I/O transformation and circuit operation. A better understanding of the I/O transformation properties of a broader neuron type/group may be necessary to define it finer granularity. In some cases, seemingly subtle variations may endow novel properties of a generic type of I/O transformation that warrant these subset of cells as belonging to a subtype. (e.g., the 15 types of retinal bipolar cells) In other cases, a continuous I/O transformation may be the essence of the circuit operation of a communication node that is achieved by implementing certain continuously variable features to its constituent cell members (e.g. orientation selective neurons in primary visual cortex - each having a particular orientation preference but the type as a whole encodes all orientations). Proper recognition of cell type granularity will also help distinguishing variations related to cell identity versus its physiological or pathological states. Therefore, the communication module concept may provide a mechanistic and functional basis to define cell type granularity that informs classification and taxonomy.

### Conclusions and perspectives

The significance of discovering and understanding neuronal diversity is now well recognized. We suggest that uncovering the biological basis of neuron type identity and granularity is necessary to decipher neuronal diversity and achieve satisfying classification and taxonomy. High throughput and large scale single cell RNAseq and epigenomic analysis will accelerate the generation of massive datasets, laying out the molecular landscape for systematic cell type discovery and classification. The revelation that each cardinal neuron type can be defined as a neural communication module with characteristic I/O transformation properties that are encoded by its key transcriptional signatures suggests that interneuron types are not just statistical constructs shaped by arbitrary criteria but are inherently biological entities built with molecular genetic mechanisms that can be understood through fundamental principles. Complementary and synergistic, these advances should together substantially clarify the work draft of interneuron taxonomy and drive cell type discovery across cerebral cortex, a strategy with broad implications in exploring cell diversity in the central nervous system.

Currently, the two basic features of neuronal communication, connectivity and I/O transformation, cannot yet be measured at scale. Technical improvement in I/O tracing and dense connectomics will facilitate the scalable measurement of neural connectivity. In parallel, continued innovation in genetic targeting, optogenetic manipulation, activity readout and perturbation will enable the characterization of I/O transformation in increasing numbers of neuronal subpopulations. Fortunately, it is increasingly evident that transcriptomes and epigenomes contain highly rich information regarding cell identity and physiological state (*62*). Indeed, transcriptome and epignenome are not just another measure of cell features, they represent *the* dataset that contains mechanistic though encrypted information of other cell phenotypes (e.g. morphology, connectivity, physiology). To retrieve and interpret such information, it is necessary not only to obtain high resolution (i.e. high depth) datasets but further to recognize and annotate “functional elements and modules” in these molecular genetic profiles. This may be best illustrated by a comparison between the transcriptomic/epigenomic approach to cell typing versus the genomic approach to species classification. In this postgenomic era, contemporary geneticists and evolutionary biologists can go a long way using genome alignment and comparative analysis to deduce the identity and evolutionary relationship of many organisms toward their classification and taxonomy. However, this approach, now almost taken for granted, can only succeed if two requirements are met. First, it is necessary to obtain high resolution (i.e. nucleotide) and high coverage (preferably full genome) sequences. Random subsets of genomes (e.g. obtained with shot-gun sequencing at 10% coverage) are far less useful for across-species genome comparison. Second, it is essential that comparative analysis of genome information is guided by genetic principles and mechanistic knowledge. Comparison of GC contents or overall genome-wide alignment are unlikely to be informative. The recognition of functional elements such as the genetic code, exons and introns, and promoters, enhancers and insulators, which constitute only a tiny fraction of the genomic landscape, is necessary to interpret and compare genome sequences, extract biologically meaningful information, and infer species identity and relationships rooted in the universal genetic information that flows through the evolutionary tree of life.

This genomic approach to species classification may apply to transcriptomic and epigenomic approaches to understanding the logic of cell type diversity in individual organisms. First, we need to obtain high resolution and high coverage omics datasets (in addition to large scale lower resolution datasets) that will provide the necessary substrates for comprehensive analysis and discovery. Second, we need to discover and apply biological principles on how genomic information is used to build a large array of molecular machines that shape cell phenotypes and properties. These may include gene batteries, cellular modules, gene regulatory networks and likely other functional elements in transcriptomes and epigenomes. Equipped with such functional and mechanistic information, biologists might be able to make significant strides in understanding cell type identity and diversity by in-depth analysis of transcriptomes and epigenomes in conjunction with phenotypic information. At a fundamental level, such an explanatory framework may facilitate understanding biological information flow from the genome to epigenomes and cell types, and to tissue organization, function and behavior, with implications in brain circuit evolution and disorders.

## Acknowledgement

We thank Drs. Brett Mensh, Adam Kepecs, Liqun Luo, and Partha Mitra for critical reading and comments of the manuscript. Z.J.H. is support by NIH U19MH114823-01, 5R01MH109665-02, 5R01MH101268-05 and the Robertson Neuroscience Fund at CSHL.

